# Non-stimulated regions in early visual cortex encode the contents of conscious visual perception

**DOI:** 10.1101/2020.11.13.381269

**Authors:** Bianca M. van Kemenade, Gregor Wilbertz, Annalena Müller, Philipp Sterzer

## Abstract

Predictions shape our perception. The theory of predictive processing poses that our brains make sense of incoming sensory input by generating predictions, which are sent back from higher to lower levels of the processing hierarchy. These predictions are based on our internal model of the world and enable inferences about the hidden causes of the sensory input data. It has been proposed that conscious perception corresponds to the currently most probable internal model of the world. Accordingly, predictions influencing conscious perception should be fed back from higher to lower levels of the processing hierarchy. Here, we used functional magnetic resonance imaging (fMRI) and multivoxel pattern analysis to show that non-stimulated regions of early visual areas contain information about the conscious perception of an ambiguous visual stimulus. These results indicate that early sensory cortices in the human brain receive predictive feedback signals that reflect the current contents of conscious perception.

## Introduction

Predictions play an important role in perception^1^. According to the theory of predictive processing, our brains use an internal model of the world to make predictions that are fed back from higher to lower levels of the processing hierarchy, thereby enabling inferences about the hidden causes of the sensory input data^2,3^. This framework might provide the key to a neuroscientific account of conscious perceptual experiences, one of the greatest challenges for theories of human brain function. Within the framework of predictive processing, it has been proposed that conscious perception corresponds to the currently most probable internal model of the world, that is, the model that makes the best predictions about the incoming sensory data^4^. From this conceptualization of conscious perception as reflecting a predictive model, it follows that predictions generated by this model should be fed back from higher to lower levels of the processing hierarchy. However, empirical studies supporting this idea are lacking. In the current study, we investigated whether predictive feedback signals that reflect the current contents of conscious perception can be observed in non-stimulated regions of human early visual cortex. Non-stimulated visual regions do not receive any bottom-up stimulation, therefore any information in these regions must come from higher visual areas through feedback connections. This approach has successfully been used in several previous studies, showing for example that feedback signals contain information not only about which visual scene is presented^5^, but also about the spatial frequency of the scene^6^. High-field fMRI studies have confirmed that decoded information in non-stimulated visual areas is due to feedback mechanisms, as this information was present in superficial cortical layers, where feedback signals arrive, and not the middle cortical layers, which process feedforward input^7^. Measuring neural activity in regions of retinotopic visual cortex that do not receive feedforward input thus provides an elegant way to isolate effects of predictive feedback signalling in the human brain. Here, we used this method to probe whether the actual contents of conscious visual perception, too, would be reflected by neural signals in non-stimulated regions of early visual cortex. We used an ambiguous motion stimulus that gives rise to bistable perception (i.e., spontaneous alternations between two perceptual states) and that was partially occluded. Decoding the two perceived visual interpretations of the constant ambiguous stimulus, rather than two distinct stimuli, from non-stimulated visual regions would thus enable us to identify the presence of feedback signals reflecting the current conscious percept.

## Results

During fMRI scanning, participants were presented with ambiguous plaid-motion stimuli, composed of two gratings moving in different directions (fig. 1A)^8^. The luminance of gratings and intersections was chosen such that the stimuli could be perceived either as two gratings moving in different directions (hereafter referred to as ‘component perception’) or as one pattern moving in the average direction of the two gratings (‘pattern perception’). We used four different stimulus configurations: The angle between the gratings could be 60° or 150°, and the average motion direction was either leftward or rightward. Crucially, one quadrant of the stimulus was always occluded, which allowed us to analyse fMRI signals in non-stimulated parts of retinotopic visual areas.

**Fig. 1.**
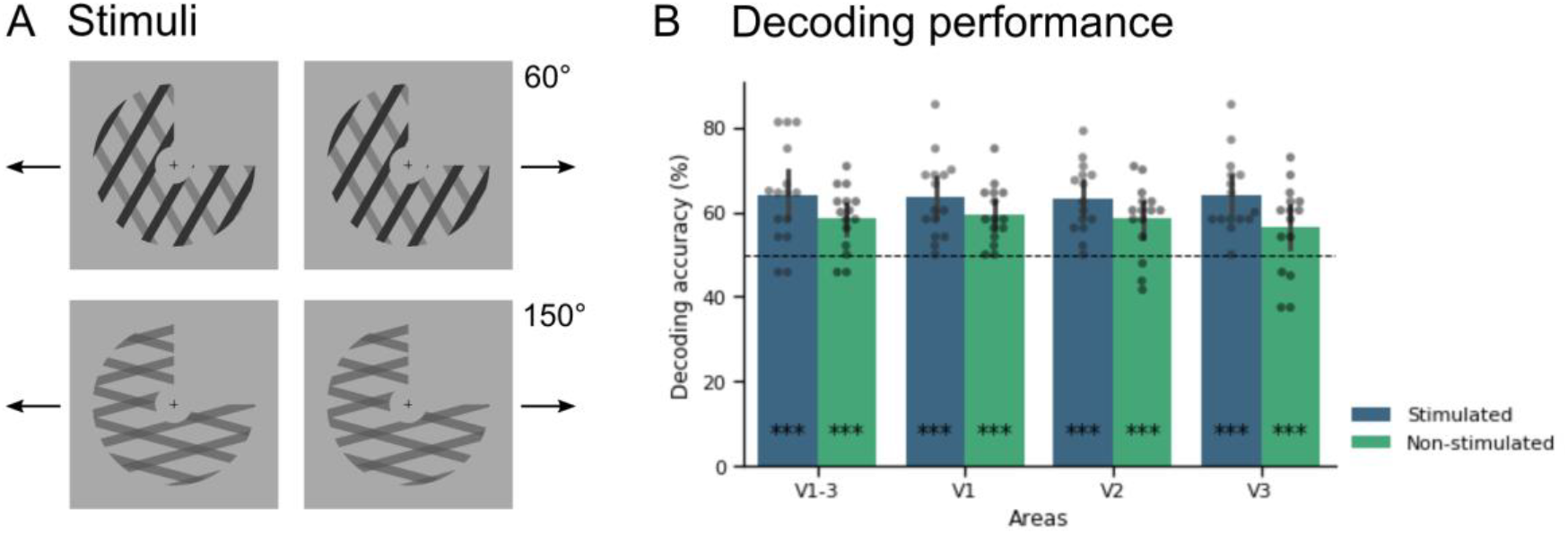
A) Ambiguous moving plaid stimuli were presented in four different stimulus configurations, which differed in the angle between the two component gratings (60° or 150°) and the overall motion direction of the resulting pattern (leftward or rightward). B) Classifier accuracy discriminating component and pattern perception across all stimulus configurations for stimulated and non-stimulated regions of early retinotopic areas. Error bars represent 95% confidence interval (CI). *p<0.05, **p<0.01, ***p<0.001.

Participants were asked to fixate the central fixation cross and indicate transitions between component and pattern percepts via button presses. Trials in which no perceptual transitions were reported were excluded. Eye tracking was performed and used for a control analysis, in which we discarded runs with poor fixation performance. Functional localisers of the stimulated area, occluded area, and border in between, as well as standard retinotopic mapping procedures, were used to delineate regions of interest for early visual regions that responded to our stimuli and for those representing the non-stimulated quadrant. For an additional control analysis, we used a V5 localiser to define hMT+/V5. We then applied multi-voxel pattern analysis using a linear support-vector-machine classifier to decode participants’ perception from both stimulated and non-stimulated regions of visual cortex for each stimulus configuration, and averaged decoding accuracies across conditions. Permutation tests were performed to determine significance (see figure S1 and Supplementary Methods for details).

As displayed in figure 1B, significant above-chance decoding performance was obtained for both stimulated (64.1%, p<0.001) and non-stimulated (58.6%, p<0.001) regions of areas V1-V3 together. Decoding performance also reached significance in each of the retinotopic areas separately (V1: 63.4% stimulated, 59.4% non-stimulated; V2: 63.3% stimulated, 58.4% non-stimulated; V3: 64% stimulated, 56.3% non-stimulated; all p<0.001). Our control analysis in which runs with poor fixation performance were discarded led to comparable results (see fig. S2 and Supplementary methods and results for details). Furthermore, when we corrected for the difference in number of voxels between our stimulated and non-stimulated regions, we still obtained significant above-chance decoding results (see fig. S3 and Supplementary methods and results for details).

According to the predictive processing theory, predictions about incoming sensory data are fed back from higher visual areas. In the case of plaid motion stimuli, area hMT+/V5 has been reported to be differentially activated during component vs. pattern motion^9^ and is therefore a likely candidate for the origin of feedback signalling. Here, we replicated the previous finding of greater hMT+/V5 activity during component motion compared to pattern motion (fig. S5). More critically, we additionally tested whether perceptual states could also be decoded from hMT+/V5 activity in a subsample of participants, as this area should be able to represent the different percepts if it feeds back predictions about these stimuli. This proof-of-concept analysis revealed that indeed the component and pattern percepts could be decoded from hMT+/V5 with high accuracy (69.0%, p < 0.001, see fig. S4 and Supplementary methods and results for details).

## Discussion

Our findings show that the current perceptual state during bistability can be decoded from fMRI signal patterns not only in stimulated early visual regions, which is in line with previous studies^10^, but crucially also in non-stimulated retinotopic visual cortex, which did not receive any bottom-up input. This suggests that non-stimulated regions of early visual cortex contain information not only about visual stimulation in the surrounding context, as previously shown^5^, but even about conscious perception independent of visual stimulation *per se*. This is in line with current theories that model bistable perception within the framework of predictive processing^4,11^. According to this view, ambiguous stimuli (such as the bistable moving plaids used here) provide equally strong sensory evidence for two different percepts, but the currently dominant percept establishes an implicit prediction regarding the cause of the sensory input. This prediction is thought to stabilize the current perceptual state through feedback from higher to lower hierarchical levels, while sensory evidence for the currently suppressed perceptual interpretation elicits prediction errors that act to destabilize the current percept, eventually leading to a perceptual change^12,13^. Here, we for the first time provide evidence supporting the notion of feedback signalling of predictions in bistable perception. Along these lines, we suggest that the percept-related information that we found in non-stimulated regions of early visual areas most likely arises from feedback signalling that originates from higher-level areas concerned with the computation of component vs. pattern motion perception, such as area hMT+/V5^9^. Our significant decoding results in hMT+/V5 support the idea that this area generates the predictions that are sent back to early visual areas, though future studies will have to provide direct causal evidence.

In conclusion, our current results provide compelling support for the notion that conscious perception reflects an internal model that generates predictions about the current state of the world, and that these predictions are fed back to the lowest levels of sensory processing to enable inferences regarding the sensory input.

## Acknowledgments

PS received support from the German Research Foundation (DFG grants STE 1430/7-1 and STE 1430/8-1).

## Author contributions

Conceptualisation, B.v.K. and P.S.; Methodology, B.v.K., G.W., and A.M.; Investigation, G.W. and A.M.; Formal analysis, B.v.K.; Writing – Original draft, B.v.K.; Writing – Review & Editing, B.v.K., G.W., A.M., and P.S.; Funding acquisition, P.S.

## Declaration of interests

The authors declare no competing interests.

## Supplementary methods

### Subjects

Sixteen participants took part in the study. Data from one participant had to be excluded, because this participant reported only one percept in certain conditions, so that the other percept of the respective condition could not be modelled (see fMRI analysis). This resulted in a final sample of 15 participants (age 18-33, M = 23.5 years, SD = 4.22, 5 male). None of the participants reported current or previous neurological or psychiatric disorders. All had normal or corrected-to-normal vision and were right-handed. Besides these general criteria, inclusion was based on performance in a previous behavioural session with the same ambiguous plaid stimuli. An average perceptual phase duration of > 4 s and a balance of at least 80/20 between the two percepts in each possible stimulus configuration (pattern and component perception, see Stimuli) were required to be selected for the fMRI session. The study was approved by the local ethics committee, and participants gave written informed consent.

### Stimuli

Plaid stimuli were created by superimposing two individual component square-wave gratings. The stimuli were designed to be perceptually ambiguous, yielding bistable perception with spontaneous alternations between perception of either the two components moving in different directions (‘component perception’) or of one pattern moving in the average direction of the two gratings (‘pattern perception’). The angle between the components could be 60° or 150°, but for both angles the average motion direction between the two gratings was horizontal, either leftward or rightward, resulting in four stimulus configurations (60° left, 60° right, 150° left, 150° right) that all elicited bistability between component and pattern perception. fMRI results were pooled across these four stimulus configurations, as they were not relevant to the purpose of the present study. The individual gratings had a spatial frequency of 0.5 cycles per degree of visual angle and a duty cycle of 0.3. The term ‘duty cycle’ refers to the proportion of the width of the darker bars within one cycle of the grating. The speed of the individual gratings was 1.3 cycles/s for the 60° stimuli, and 0.39 cycles/s for the 150° stimuli. The speed of the resulting plaid stimuli was 1.5 cycles/s for all stimulus configurations.

The plaid stimuli were presented within a centred annulus with a diameter of 13° of visual angle. In the centre of the annulus, which had a diameter of 3°, a fixation cross was presented. The background surrounding the stimuli had a luminance of 40 cd/m^2^. The luminance of the gratings of the 150° stimuli was 14 cd/m^2^. For the 60° stimuli, the two component gratings differed in luminance: one grating had 2 cd/m^2^, the other 20 cd/m^2^. The luminance of the intersections of the gratings was determined in pilot experiments that aimed at approximate equiprobability of component and pattern perception for all stimulus types and resulted in an intersection luminance of 9 cd/m^2^ for the 150° stimuli and 2 cd/m^2^ for the 60° stimuli.

### Procedure

The stimuli were presented on a screen at the end of the MRI scanner bore. Participants laid in the scanner in supine position and viewed the stimuli on the screen through an angled mirror. They were asked to fixate on the central fixation cross and report their percept (pattern or component perception) by button presses. They had to report their percept as soon as the stimulus was presented, and press a button anytime their percept changed. A pattern percept was reported with the right index finger, and a component percept with the right middle finger. Each run comprised eight trials, lasting 60 s each, during which a plaid stimulus was continuously presented in one of the four stimulus configurations. Each trial was followed by a 10 s fixation interval, during which only the fixation cross was presented. Each stimulus configuration was presented twice per run in pseudorandomised order. There were six runs in total.

After the main experiment, two functional localisers were presented. The first was a stimulus localiser. Here, each stimulus from the main experiment was presented for 12 s, followed by fixation for 8 s, in a block-design. Different from the main experiment, participants were asked to fixate only and not report their perception. All conditions were presented four times in total. This functional stimulus localiser allowed for selection of voxels that were activated by the stimuli used in the main experiment. Furthermore, we used a functional localiser that mapped the non-stimulated region and was designed to preclude any spill-over of activity from the stimulated region, similar to the localiser of Smith & Muckli (2010). During this localiser, participants viewed contrast-reversing checkerboard stimuli (4Hz), which were again presented for 12 s each, followed by 8 s of fixation. Each condition was repeated 8 times. The localiser contained ‘surround stimuli’, mapping the border between stimulated and non-stimulated regions, and ‘target stimuli’, mapping the non-stimulated region. The surround stimulus was presented at 0.5° of visual angle diagonally from the fixation cross, mapping the outer 1° of the non-stimulated quadrant (see figure S1A). The checkerboard representing the non-stimulated quadrant, i.e. the target stimulus, was presented at 1° diagonally from the surround stimulus (see figure S1B). Thus, the target region, from which voxels were selected for our decoding analysis of the non-stimulated quadrant, was ~2° away from the stimulated region. The scanning session ended with a structural T1 scan (MPRAGE). Standard phase-encoded retinotopic mapping was performed in a separate scanning session to define regions V1-3.

**Fig. S1.**
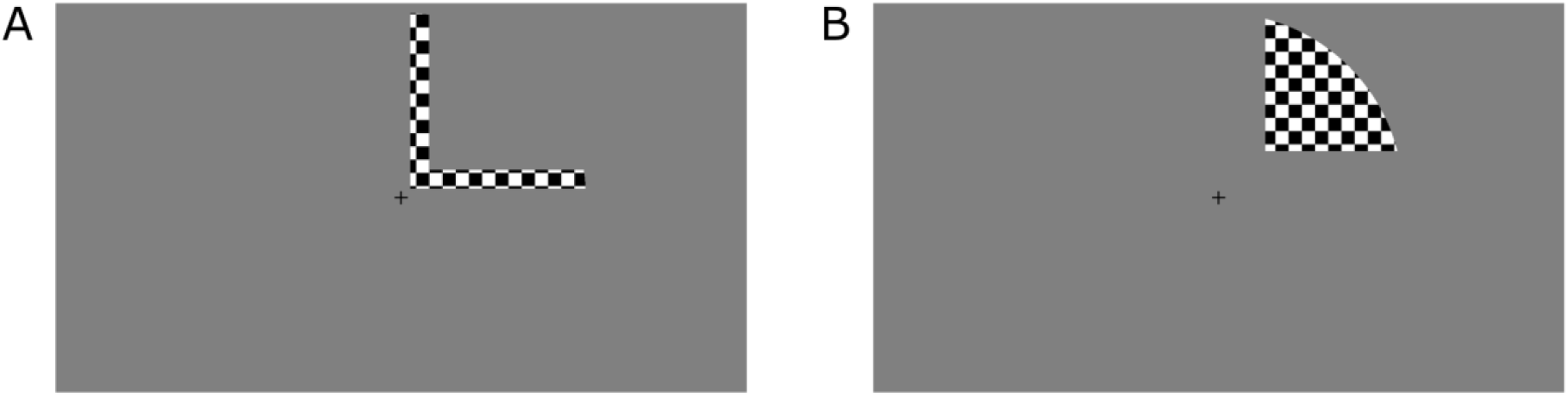
A) The surround stimulus mapping the border between stimulated and non-stimulated regions. B) The target stimulus mapping the non-stimulated quadrant.

### Scanning parameters

Functional MRI data were acquired using a 3 T TIM Trio scanner (Siemens, Erlangen, Germany), equipped with a 12-channel head-coil. A gradient echo EPI sequence was used (TR: 2 sec, TE: 30 msec, flip angle: 78°, voxel size 2.3 × 2.3 × 2.3 mm). Slices were oriented parallel to the calcarine sulcus and acquired in descending order. A total of 135 volumes were acquired for each run of the main experiment, 163 volumes for the stimulus localiser, 163 volumes for the non-stimulated quadrant localiser, 123 volumes per run (3 in total) for the polar angle retinotopic mapping, and 102 volumes per run (3 in total) for eccentricity mapping. Anatomical images were obtained using an MPRAGE sequence (TR: 1.9 sec, TE: 2.52 msec, flip angle: 9°).

### Eye movements

Eye movements were recorded with an iView Xtm MRI-LR system [SensoMotoric Instruments (SMI), Teltow, Germany] using a sampling rate of 50 Hz. Due to technical difficulties, no usable eye tracking data were obtained for four participants, and for one run of a fifth participant. The eye tracking data were used in a control analysis to discard runs with poor fixation performance. To determine fixation performance, a radius of 1.5° from fixation was defined as the fixation area. Eye movements beyond this area were considered as outliers. Data were detrended and mean-corrected to determine the number of these outliers, and runs in which eye movements extended beyond 1.5° of fixation in more than 5% of all data points were excluded. A total of 10 runs distributed across 5 participants were excluded in the control analysis based on eye tracking exclusion criteria.

### fMRI analysis

The fMRI data were preprocessed and analysed using SPM12. First, the functional images were realigned to correct for head motion, after which they were coregistered with the structural image obtained in the same session. Then, both functional and structural images were coregistered with the structural image obtained in the retinotopy session. No normalisation or smoothing was applied, as is common for studies using MVPA.

A general linear model (GLM) was set up with regressors modelling the participants’ percepts (pattern vs components) of each condition, resulting in eight regressors of interest. Motion parameters as well as a regressor modelling fixation in between trials were included as regressors of no interest. If participants reported only one percept for a certain condition, the other percept of that condition could not be modelled in that run; therefore, such runs were excluded. This affected all runs from one participant, and another 7 runs distributed across 3 participants.

### ROI definition

Regions of interest (ROIs) were defined with similar methods as those used by Smith & Muckli (2010). First, regions V1-V3 were defined using standard retinotopic mapping procedures. Within regions V1-3, only the voxels that showed significant positive response to the stimulated region (t-contrast stimulus > fixation, p < 0.01 uncorr.) in our stimulus localiser were selected. For the non-stimulated region, the following procedure was used. First, only voxels that showed significant positive response to the target region (t-contrast stimulus > fixation, p < 0.01 uncorr.) were selected. Then, in order to ensure that these voxels were not also responsive to the stimulated region, we further selected only the voxels that met these criteria: significant positive response to the non-stimulated target area alone (t > 1.65, p < 0.01 uncorr.), no significant response to the stimulated area alone (t > 1.65, p < 0.01 uncorr.), and no significant response to the surround region (t > 1.65, p < 0.01 uncorr.).

The stimulated ROIs were naturally larger than the non-stimulated ROIs, as the stimulus spanned three quadrants compared to one occluded quadrant. Furthermore, our strict criteria for selecting non-stimulated voxels outlined above meant we only selected a small sample of the voxels corresponding to the occluded quadrant. To correct for potential biases induced by this difference in ROI size, we performed an additional control analysis with smaller stimulated ROIs that had the same number of voxels as their non-stimulated counterpart ROI. These ROIs were generated by manually selecting voxels corresponding to the stimulus quadrant immediately opposite the occluded quadrant, in our case the quadrant in the upper left visual field. As such, we selected voxels in the right hemisphere below the calcarine sulcus. From these voxels, we randomly selected *n* voxels, with *n* being the number of voxels of the non-stimulated ROI for that particular visual area (V1-3) and participant. For two participants, not enough voxels were available in the respective stimulated quadrant of V1 to match the number of voxels from the non-stimulated V1 ROI. For these two participants, we therefore used all the voxels available in the stimulated quadrant and thus had slightly less voxels in stimulated V1 ROI compared to the non-stimulated V1 ROI (for one participant 12 stimulated voxels vs 15 non-stimulated voxels, for the other participant 6 stimulated voxels vs 24 non-stimulated voxels).

### MVPA

Multi-voxel pattern analysis (MVPA) was performed using The Decoding Toolbox (Hebart, Görgen, & Haynes, 2015), which implements LibSVM software (http://www.csie.ntu.edu.tw/wcjlin/libsv). A linear support vector machine was trained to discriminate pattern from component percepts based on the beta values resulting from the GLM. This classification was performed for each stimulus configuration separately. Classifier performance was tested using a leave-one-run-out cross-validation approach. Training was carried out on all but one run, which served as the test data. This was repeated until all runs had served as a test run once. The decoding accuracy was averaged across cross-validations and then across conditions. Permutation testing was conducted to determine the significance at the group level as described by Stelzer, Chen, & Turner (2013). In brief, we provided the classifier with all possible combinations of shuffled label assignments for each participant and performed the decoding procedure for each label assignment. Then, we randomly selected one of these decoding accuracies from each participant and calculated the mean decoding accuracy. This procedure of random selection and calculation of mean decoding accuracy was repeated 10,000 to generate a distribution of decoding accuracies. We then used a cut-off of 95% to determine significance of our results.

## Supplementary results

### Phase durations

The mean perceptual phase duration of the 60° stimuli (averaged across leftward and rightward moving stimuli) was 7.4 s for components (SD = 8.6) and 9.9 s for patterns (SD = 4.6). For the 150° stimuli, mean phase duration for components was 8.2 s (SD = 7.5) and for patterns 4.9 s (SD = 1.7).

### Control analysis discarding runs with poor fixation performance

Overall fixation accuracy across all participants was 97.3%. Despite this high accuracy, we performed a control analysis discarding runs with fixations more than 5% outside of our fixation ROI. As displayed in figure S2, significant above-chance decoding performance was obtained for both stimulated (64.0%, p<0.001) and non-stimulated (58.9%, p<0.001) regions of areas V1-V3 together. Decoding performance also reached significance in each of the retinotopic areas separately (V1: 62.9% stimulated, p<0.001, 57.8% non-stimulated, p=0.015; V2: 62.4% stimulated, p<0.001, 58.0% non-stimulated, p = 0.007; V3: 63.0% stimulated, p<0.001, 56.7% non-stimulated, p<0.001).

**Fig. S2.**
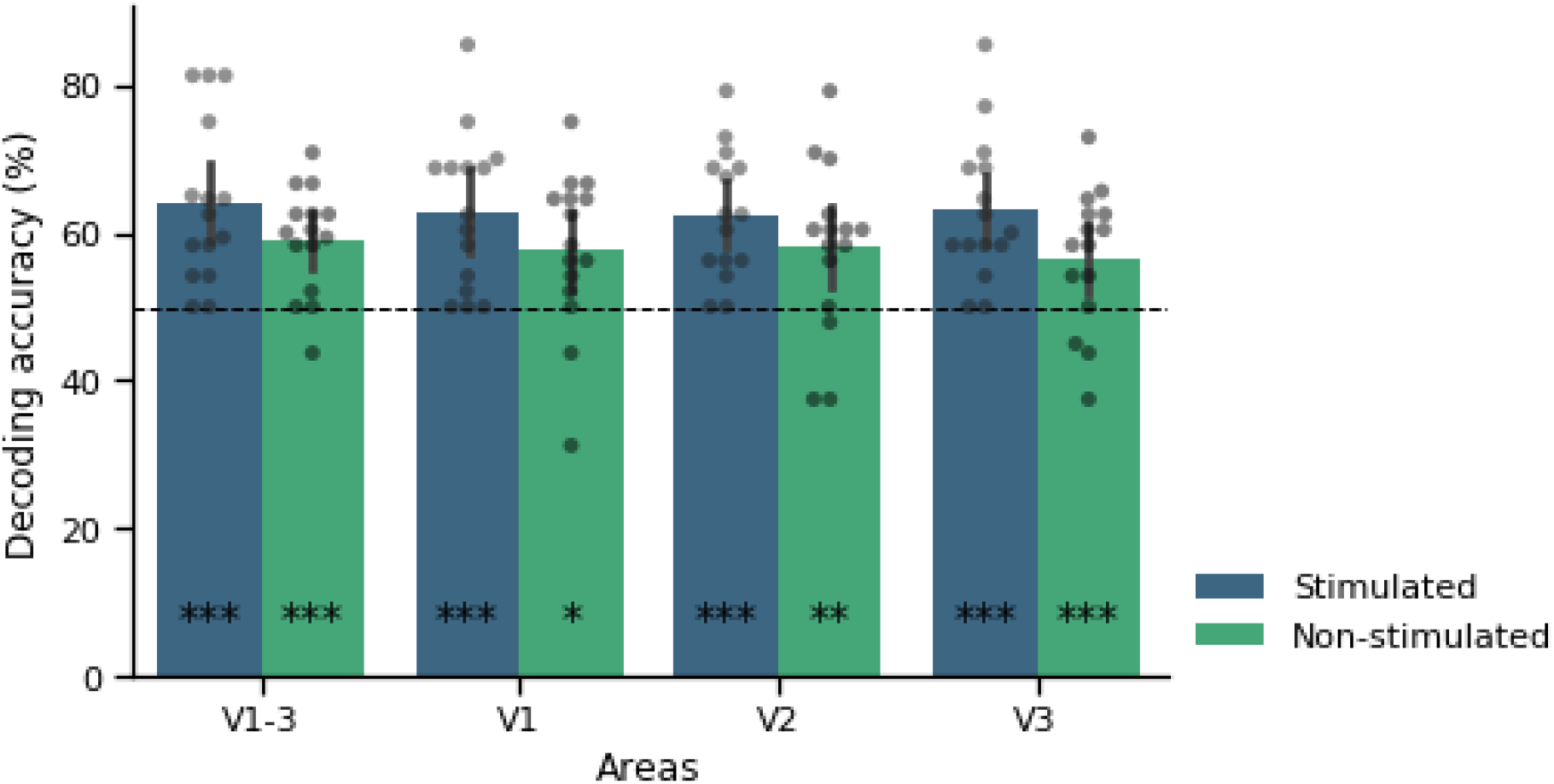
Classifier accuracy discriminating component and pattern perception across all stimulus configurations for stimulated and non-stimulated regions of early retinotopic areas. In this analysis, runs with poor fixation performance were excluded. Error bars represent 95% confidence interval (CI). *p<0.05, **p<0.01, ***p<0.001.

### Control analysis correcting for the difference in number of voxels between stimulated and non-stimulated ROIs

In this analysis, we decoded from stimulated and non-stimulated ROIs that were matched in size. As displayed in figure S2, significant above-chance decoding performance was obtained for both stimulated (60.9%, p<0.001) and non-stimulated (58.6%, p<0.001) regions of areas V1-V3 together. Decoding performance also reached significance in each of the retinotopic areas separately (V1: 55.2% stimulated, 59.4% non-stimulated; V2: 56.5% stimulated, 58.4% non-stimulated; V3: 59.2% stimulated, 56.3% non-stimulated, all p<0.001).

**Fig. S3.**
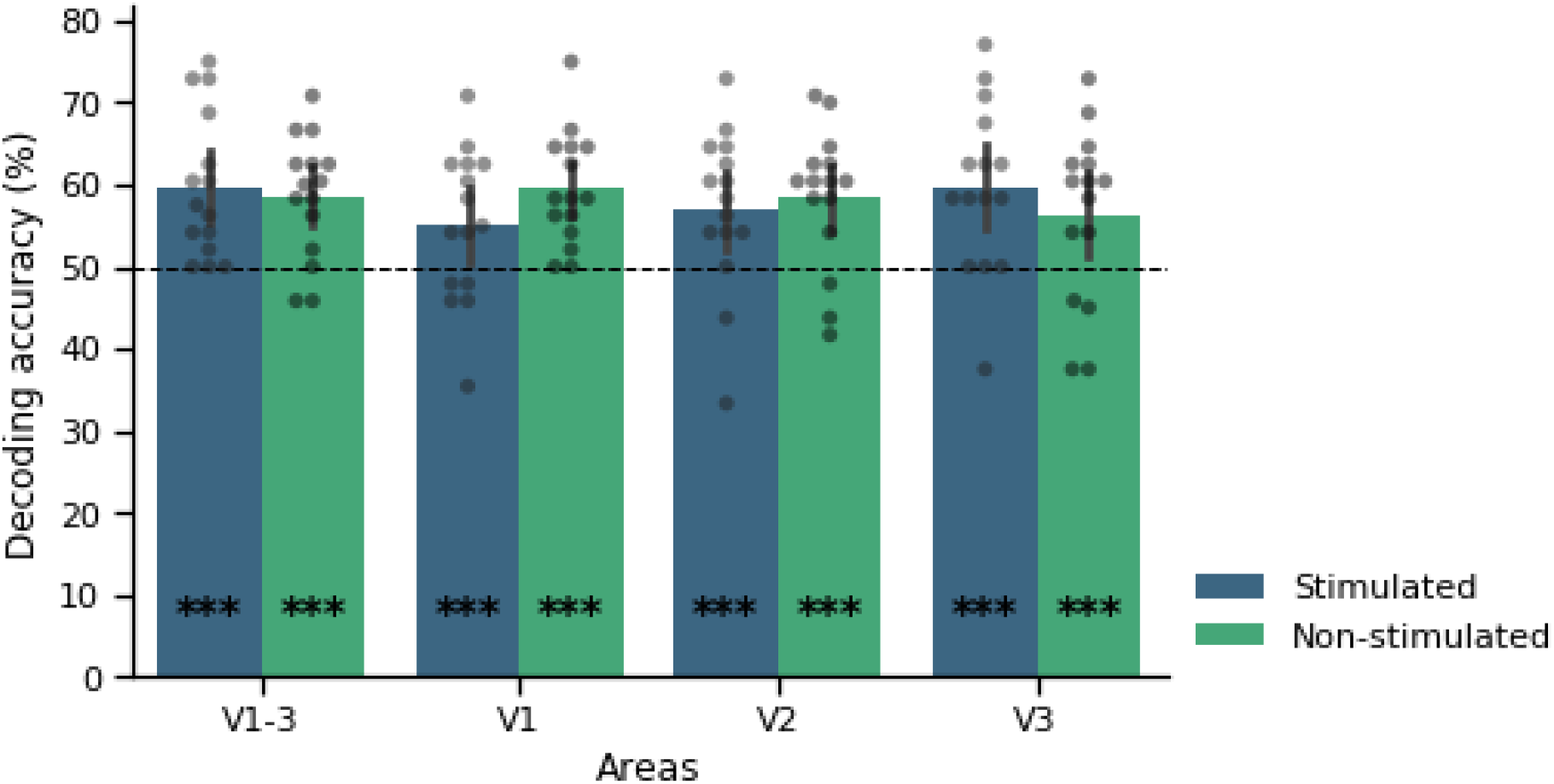
Classifier accuracy discriminating component and pattern perception across all stimulus configurations for stimulated and non-stimulated regions of early retinotopic areas. In this analysis, the number of voxels in stimulated V1 ROIs matched those of non-stimulated V1 ROIs. Error bars represent 95% confidence interval (CI). *p<0.05, **p<0.01, ***p<0.001.

### Control analysis decoding from hMT+/V5

hMT+/V5 localiser data were available for 10 of our subjects. From these hMT+/V5 ROIs, we could decode component vs pattern percepts significantly above chance (69.0%, p < 0.001; see figure S4).

**Fig. S4.**
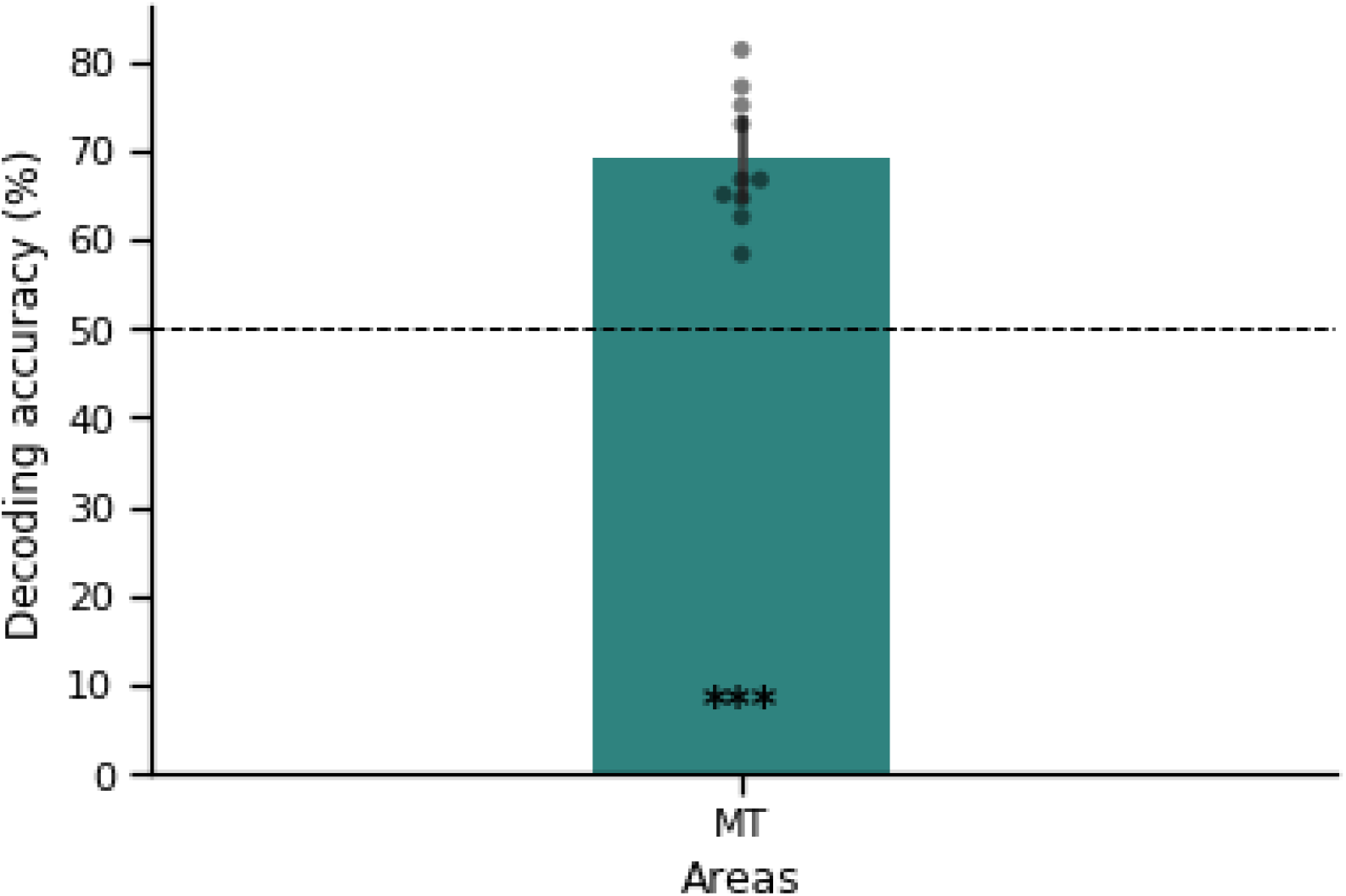
Classifier accuracy discriminating component and pattern perception across all stimulus configurations for area hMT+/V5. Error bars represent 95% confidence interval (CI). *p<0.05, **p<0.01, ***p<0.001.

### Control analysis testing for non-specific effects in early visual cortex

In order to test whether non-specific effects related to the change in perception and resulting decision making influenced our results, we performed a univariate analysis contrasting component with pattern percepts and vice versa. To this end, data preprocessing included coregistration of functional and anatomical images, normalisation to MNI space and smoothing with an 8mm full width at half maximum kernel. The same GLM was run as was used for our MVPA analysis. T-contrasts of components > patterns and patterns > components were passed on to group level T-tests. An initial voxel threshold of p < 0.001 uncorrected was used with FWE cluster correction to determine significance.

Since it has been shown that components elicit more activity in hMT+/V5 than patterns (Castelo-Branco et al., 2002), we expected clusters in hMT+/V5 for the contrast components > patterns. As such we performed a ROI analysis using an anatomical mask of hMT+/V5 from the anatomy toolbox. This contrast indeed revealed clusters in bilateral hMT+/V5, supporting the results by Castelo-Branco et al. (2002). No other clusters reached significance. The reverse contrast, patterns > components, also yielded no significant clusters. These results suggest that no non-specific effects influenced our decoding results in visual cortex.

**Fig. S5.**
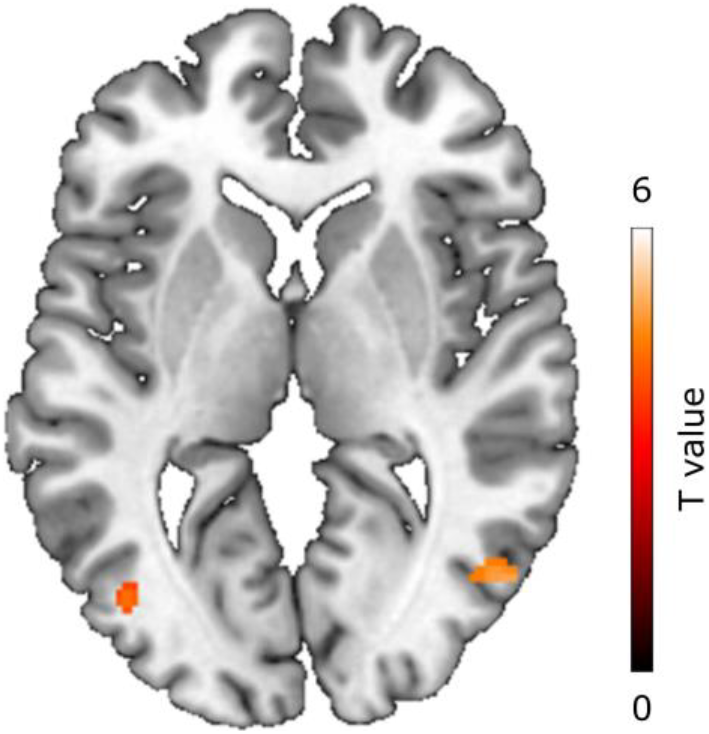
Univariate analysis showing increased activity for components compared to patterns in bilateral hMT+/V5. ROI analysis with anatomical hMT+/V5 ROI from the anatomy toolbox using an initial voxel threshold of p < 0.001, uncorrected, showing FWE cluster corrected results.

## Notes

### Competing Interest Statement

The authors have declared no competing interest.

